# Modeling Bistable Dynamics Arising from Macrophage-Tumor Interactions in the Tumor Microenvironment

**DOI:** 10.1101/2025.06.24.661275

**Authors:** Hwayeon Ryu, Susanna Röblitz, Kamila Larripa, Anna-Simone Frank

## Abstract

Macrophages in the tumor microenvironment (TME), known as tumor-associated macrophages (TAMs), originate primarily from circulating monocytes that differentiate under the influence of tumor-derived signals. Within the TME, naïve macrophages can adopt either a pro-inflammatory, anti-tumor (M1-like) or anti-inflammatory, pro-tumor (M2-like) phenotype. These phenotypic shifts significantly affect tumor progression, making TAMs attractive targets for therapeutic intervention aimed at blocking recruitment, promoting anti-tumor polarization, or disrupting tumor–macrophage interactions. In this study, we develop a mathematical model capturing the temporal dynamics of tumor volume alongside populations of naïve, M1-like, M2-like, and mixed (M1/M2) phenotype TAMs. The model incorporates the bidirectional influence between tumor development and macrophage polarization. We conduct the bifurcation as well as global sensitivity analyses to identify regions of bistability for tumor dynamics in the parameter space and the impact of sensitive parameters on TME. The model results are then linked to treatment strategies that may effectively induce transitions from high to low tumor burden.

**Highlights:**

- We propose a mathematical model to describe the temporal evolution of the tumor volume alongside different phenotypes of tumor-associated macrophages (TAMs).
- We investigate the impact of TAMs on tumor growth and decline as well as the influence of the tumor on transition rates between different TAM phenotypes.
- Bifurcation and global sensitivity analyses reveal the regions of bistability for tumor dynamics and the impact of sensitive model parameters on the tumor microenvironment.
- Model results are linked to treatment strategies that manipulate the bistable system to transition from high to low tumor volume.

## 1. Introduction

The immune system is a crucial part of the tumor microenvironment (TME), which consist of cancer cells, stromal tissue and extracellular matrix [1]. Among immune cells in the TME, macrophages, also referred to as tumor associated macrophages (TAMs), represent the most abundant population type [2]. Within the TME, they are primarily derived from circulating monocytes [2]. Based on external stimuli, TAMs can acquire different phenotypes, which are broadly classified into pro-inflammatory (M1) or anti-inflammatory states (M2) [2]. They can also switch their phenotype in a process called polarization. In cancer, the M1 and M2 polarized states have contrasting effects on tumor growth and progression, where M1 is tumor killing and M2 tumor promoting [3].

TAMs, in particular, are a potential target for cancer therapies because they represent a major proportion of myeloid cells in a solid tumor. The elimination of tumor cells by the immune system is based on the capacity of M1 marophages to inhibit tumor growth and spread [4]. However, this works only to some extent, more precisely, until tumor cells change their presented antigens to escape the immune surveillance leading to a tumor promoting microenvironment supported by immune cells [1]. The escaping process is associated with a higher amount of M2 phenotypes which is caused by change in cytokine levels in the TME [5]. A high ratio of M2/M1 phenotypes is currently considered a diagnostic marker for a poor outcome [2]. More generally, the distribution of different macrophage phenotypes has been linked to the prognosis of various cancers (e.g., small cell lung cancer, colorectal cancer, ovarian cancer, pancreatic cancer, etc.) [6, 7, 8, 9]. Vice versa, tumor size impacts the immune system composition [10], resulting in a subtle interplay between the immune system and tumors as they grow.

Developing mathematical models of macrophage polarization in TME is crucial to better understand the complex processes involved in the activation and function of macrophages, including their role in disease. These models help validate hypotheses, make predictions, and can be used to analyze how different factors influence macrophage polarization, ultimately aiding in the development of more effective therapies. Many mathematical models have been developed to study the interactions between tumor cells and TAMs [11, 12, 11, 13]. These models typically focus on inter-cellular signaling pathways on a single-cell level or on the temporal development of cell populations. Besides agent-based [14, 15] and spatio-temporal models [16], the majority of cell population models use an ordinary differential equations (ODE) framework, whereby the state variables represent different cell types, including one [17], two (M1 and M2) [18, 19, 20, 21] or three (M1, M2, and mixed) [22, 23] macrophage phenotypes. These models focus on studying the impact of tumor decomposition in terms of cell types on its development [18, 22, 23, 24] or on understanding the process of repolarization from M1-like to M2-like macrophages [20]. Despite the rich results of previous studies on macrophage polarization in TME, many models previously developed exhibit only mono-stability in tumor–macrophage dynamics.

However, tumors are inherently bistable systems [25]. Common regulatory motifs reported for macrophage transitions, such as positive or double-negative feedback loops, exhibit a switch-like behavior, suggesting the potential for bistability of the system, and there is evidence for multistability (including bistability) in macrophage activation pathways at different molecular levels. Multistability has been studied in ODE models for cellular signaling pathways on a single-cell level [26, 14, 27] and in a PDE-model based on a continuous phenotype spectrum [28], with both resolving and chronic inflammatory steady states permissible, but to our knowledge bistability has so far not been studied in ODE models on a cell-population level.

Many immunotherapies specifically target macrophages [29, 30]. This includes increasing the death rates of M2-like macrophages, reducing the recruitment of monocytes and proliferation of macrophages, the repolarization of M2-like macrophages, and stimulating the killing of tumor cells. Treatments include a broad array of approaches, such as monoclonal antibodies [31], immune checkpoint inhibitors [32], administration of cytokines (e.g., interleukin and interferon) [33], immunomodulators which either increase or decrease the production of antibodies [34], and chimeric antigen receptor (CAR) T-cell therapies [35] (a type of gene therapy).

Hence, the aim of our study is to model bistable dynamics arising from macrophage–tumor interactions in TME by identifying conditions that lead to bistable states, and explore their implications for immunotherapy strategies using simulation-based analysis. For this purpose, we develop the model adapted and modified from [22] by additionally including naïve macrophages into our model due to their critical role in initiating the process of macrophage polarization [36]. In addition, inspired by our prior work on a stochastic model of macrophage polarization [37], we include direct transitions from M1-like to M2-like macrophages that do not involve the mixed phenotype. We perform parameter perturbation studies and identify regions of bistability on our extended model, which we relate to potential therapeutic interventions (e.g., immunotherapies). We further perform the global sensitivity analysis (GSA) and, based on the GSA results, assess the impact of sensitive model parameters on TME.

Our paper is structured as follows: Section 2 provides the description of mathematical model we develop (Sec. 2.1), the necessary conditions for the existence of bistability (Sec. 2.2), numerical methods we employ for bifurcation analysis as well as GSA (Sec. 2.3). Section 3 describes our results for bistability analysis (Sec. 3.1), for GSA (Sec. 3.2), and the impact of sensitive parameters based on bifurcation analysis (Sec. 3.3). We discuss our main findings in a broader context in Section 4.

## 2. Mathematical Model and Methods

### 2.1. Mathematical model and baseline parameters

Our model is adapted and modified from [22], where the authors considered the dynamics of tumor volume (*T*), M1-like macrophages (*M*_1_), M2-like macrophages (*M*_2_), and mixed (a combination of *M*_1_and*M*_2_) phenotype macrophages (*M*_*m*_). We additionally include naïve macrophages (*M*_0_) into our model due to their critical role in initiating the process of macrophage polarization [36]. We assume that *M*_0_ macrophages can differentiate into *M*_1_ or *M*_2_. Note that the model presented here only considers monocyte-derived macrophages but not tissue-resident macrophages, which have other origin and functions. Note that, for simplicity, we merge the recruitment of monocytes and their differentiation into naïve macrophages into a single rate c (Figure 1).

**Figure 1:**
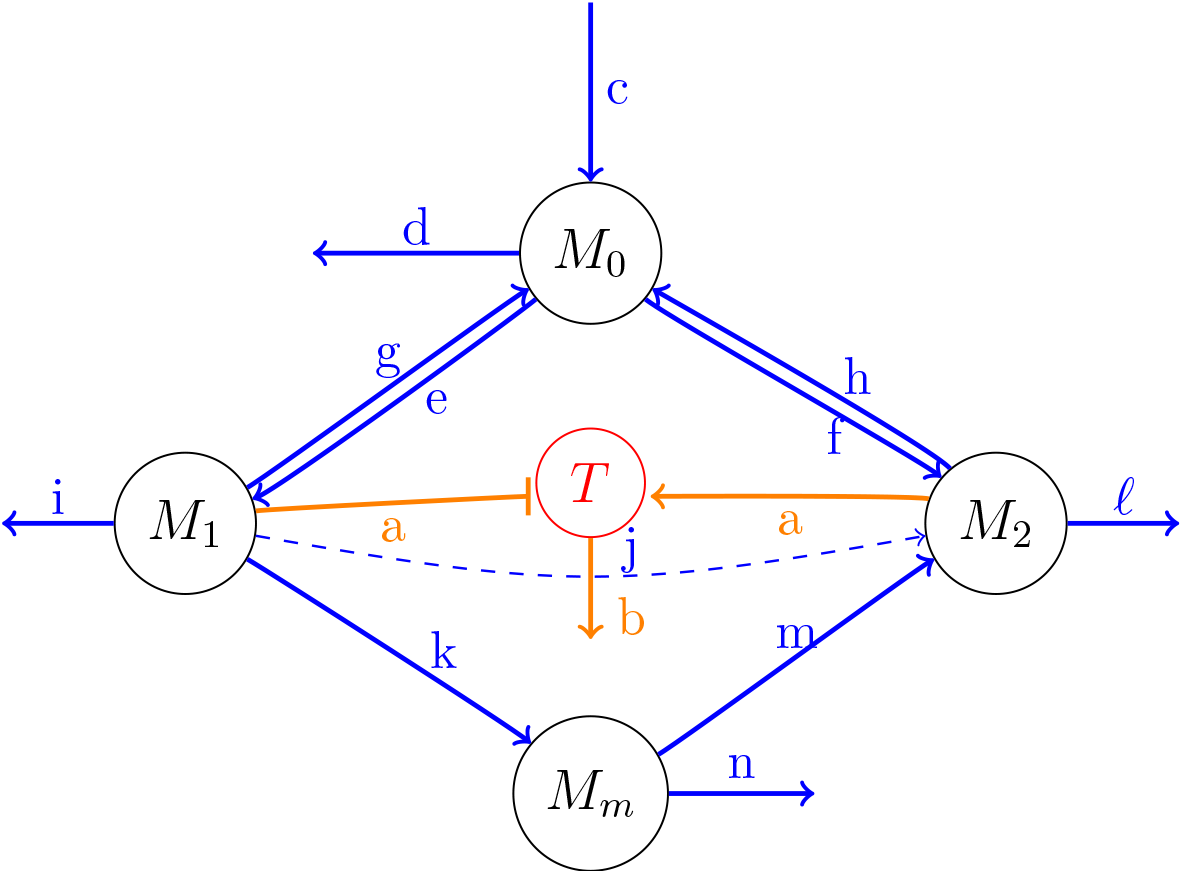
Flow-diagram of mathematical model in Eqs. (1)–(5). For a notation of biological pathways, see Table A.4 in the appendix. The dashed arrow indicates a rate that is set to zero in the baseline parameter setting but non-zero case was considered in the subsequent analysis.

The rates of transitions are captured in the parameters α_*ij*_ where i indicates the starting phenotype and j represents the transitioned phenotype (e.g., α_01_ is the rate of naïve macrophages transitioning to M1-like macrophages). Eftemie et al. [22] modeled reversible phenotype transitions between *M*_1_ and *M*_*m*_ as well as between *M*_2_ and *M*_*m*_, but they did not consider direct transitions between *M*_1_ and *M*_2_. We decide to include the transition from M1 to M2 in our analysis because polarization towards the M2-like phenotype does not always occur after a mixed or naive state but can be induced directly by specific stimuli like IL-4 and IL-13 [38].

The model equations are as follows:

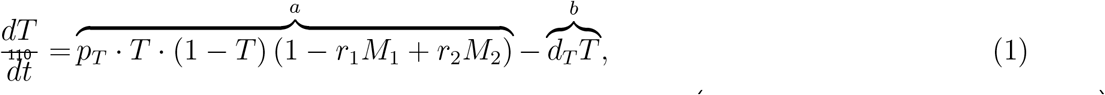

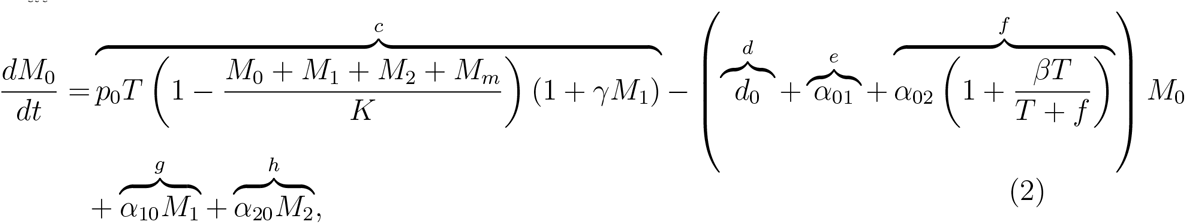

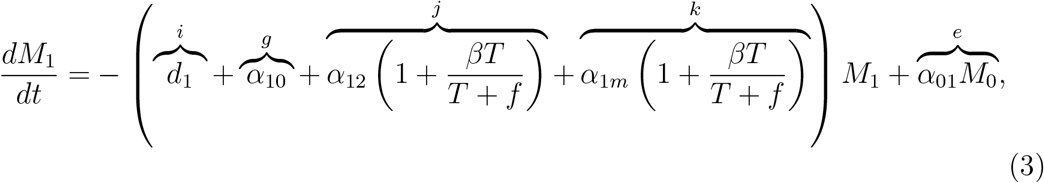

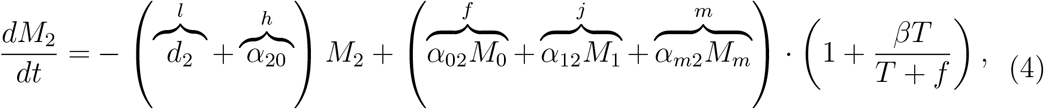

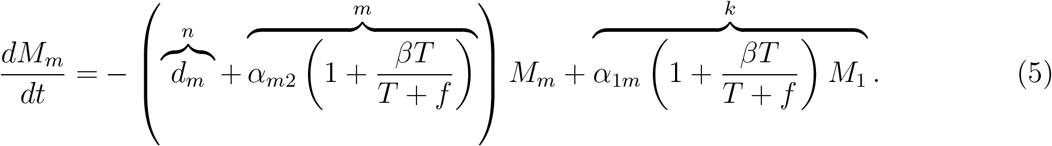

For a notation of biological pathways and their corresponding descriptions (a–n appeared in Eqs. (1)–(5) as well as in Fig. 1), see Table A.4 in the appendix. In contrast to [22], we model tumor growth logistically, which, hence, implies *T* ∈ [0, 1]. Furthermore, we model the influence of different macrophage phenotypes on tumor growth and decline differently from [22]. Since Eq. (1) represents a net effect between cell division and cell death, we decide to include an effect of both *M*_1_ and *M*_2_ on the term for cell division [39, 40]. An additional motivation to modify the model from [22] is the occurrence of steady states in their model, in which M1-like or M2-like macrophages go extinct despite large volume (Fig. 9(i) and (ii) in [22]), which is not physiologically meaningful.

To keep our model simple, we do not include any effect of the mixed phenotype on the tumor volume. While macrophages can directly induce apoptosis in tumor cells, apoptosis can also occur independently of macrophage involvement due to intrinsic mechanisms within the tumor cells themselves [41]. Hence, we add a term for macrophage-independent cell death (*d*_*T*_) in Eq. (1). This formulation allows us to derive a direct relationship between the steady-state value of *T*(*T*^*ss*^) and those of 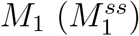 and 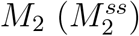, and the expression of non-trivial steady-state solution *T*^*ss*^:

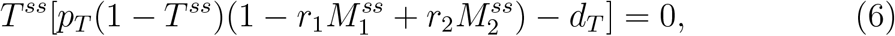

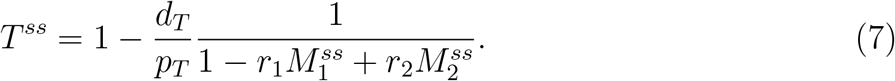

We will use the above formulation to check for the existence of multiple stable steady states in Sec. 2.2.

Circulating monocytes are recruited from the bloodstream to the site of the tumor, where they are differentiated into naïve macrophages [42]. In our model, the basal migration rate p_0_ in Eq. (2) requires the presence of tumor cells *T*and is amplified by the presence of *M*_1_ in an additive fashion since both cell types secrete signals which increase migration. For example, tumors can secrete the chemokine CCL5 and cytokines which recruit monocytes [43]. Similarly, M1-like macrophages can secrete proinflammatory cytokines TNF-α, IL-1*β*, IL-6, IL-12, IL-23 [44]. Moreover, inflammatory monocytes express CCR2, a receptor for CCL2 [45], which is secreted by both TAMs and tumor cells and induces chemotaxis in monocytes [46, 47, 48]. We limit the basal migration rate p_0_ by a carrying capacity K which reflects the portion of the tumor volume that can be colonized by different types of macrophages. Furthermore, the transition from M1-like macrophages to a naïve phenotype (captured by α_10_ in Eq. (2)) and an analogous transition from M2-like (denoted by α_20_) have been shown experimentally in vitro when the proinflammatory cytokine signal (e.g., IFN-γ or IL-4) is removed [49, 50]. Tumor cells often secrete TGF-*β*, which promotes TAMs polarization to an M2-versus-M1 phenotype. This further promotes TGF-*β* production, thus closing a positive feedback loop [51]. Hence, similar to [22], we model that tumor cells promote polarization from *M*_1_ to *M*_*m*_ (with rate of α_1*m*_) and from *M*_*m*_ to *M*_2_ (α_*m*2_) because tumor cells secrete cytokines along with other factors, which directly promote the polarization towards M2-like macrophages [39]. In addition, we account for tumor-independent polarization through the term 1 + *βT*/(*T*+ *f*) in Eqs. (3)–(5) (pathways k and m). In addition, we include this term on the rates α_02_ and α_12_, which represent polarization towards the M2-like phenotype but were not considered in [22].

The description of model parameters and their baseline values are provided in Table 1. For the proliferation rate of tumor cells (*p*_*T*_ = 0.023), we adopt the value from [22]. For the proliferation rate of naïve macrophages (*p*_0_ = 0.7) we use the same value that Eftimie et al. [22] used as proliferation rates for *M*_1_, *M*_2_, and *M*_*m*_. Since TAMs form approximately 50% of tumor volume [52] and tumor volume in our model is bounded by 1, we assume that the recruitment rate is bounded by a carrying capacity K = 0.5. Similarly, we assume the tumor to start being *M*_2_-promoting when it reaches a relative volume of f = 0.5. In this case (*T*= *f*), the transition rates towards *M*_2_ are set to be doubled by choosing *β* = 2. We also assume equal death rate for all four types of macrophages and adopt the value for M2-like macrophages from [22] (*d*_0_ = *d*_1_ = *d*_*m*_ = *d*_2_ = 0.1). Since the tumor killing rate in our model does not depend on macrophages (unlike [22]), we decide to fix it at a value (*d*_*T*_ = 0.01) that is one order of magnitude lower than the value in [22] in order to keep the whole decay rate term in the same range of value as in [22]. Since the steady state values of *M*_1_ and *M*_2_ are bounded from above by *K* = 0.5, we decide to use a value for *r*_1_ and *r*_2_ that is one order of magnitude larger (*r*_1_ = *r*_2_ = 10), as this value acts as an amplifier for the difference between *M*_1_ and *M*_2_ in steady state. We fix the remaining parameters at the values given in Table 1 but allow them to vary for the purpose of sensitivity and bifurcation analysis (see Sec. 3.2 and Sec. 3.3).

**Table 1:** Parameter description and values in the baseline setting (case 1). The marker * indicates an initial guess for a parameter value that was then varied in subsequent sensitivity and bifurcation analysis.

| Parameter | Description | Unit | Value | Reference |
| --- | --- | --- | --- | --- |
| $p_T$ | Proliferation rate of tumor cells, $T$ | $\frac{1}{\text{day}}$ | 0.023 | [22] |
| $p_0$ | recruitment rate of naïve macrophages, $M_0$ | $\frac{1}{\text{day}}$ | 0.7 | [22] |
| $K$ | maximum amount of macrophages relative to tumor volume | — | 0.5 | [52] |
| $f$ | relative required tumor size to support $M_2$ polarization | — | 0.5 | * |
| $\gamma$ | contribution of $M_1$ to recruitment of $M_0$ | $\frac{1}{\text{day}}$ | 1 | * |
| $r_1$ | contribution of $M_1$ to the killing of tumor cells | $\frac{1}{\text{cells}}$ | 10 | * |
| $r_2$ | contribution of $M_2$ to the proliferation of tumor cells | $\frac{1}{\text{cells}}$ | 10 | * |
| $d_T$ | apoptosis rate of tumor cells | $\frac{1}{\text{day}}$ | 0.01 | modified from [22] |
| $d_0$ | apoptosis rate of $M_0$ | $\frac{1}{\text{day}}$ | 0.1 | modified from [22] |
| $d_1$ | apoptosis rate of $M_1$ | $\frac{1}{\text{day}}$ | 0.1 | modified from [22] |
| $d_2$ | apoptosis rate of $M_2$ | $\frac{1}{\text{day}}$ | 0.1 | modified from [22] |
| $d_m$ | apoptosis rate of $M_m$ | $\frac{1}{\text{day}}$ | 0.1 | modified from [22] |
| $\alpha_{01}$ | transition rate from $M_0$ to $M_1$ | $\frac{1}{\text{day}}$ | 1 | * |
| $\alpha_{02}$ | transition rate from $M_0$ to $M_2$ | $\frac{1}{\text{day}}$ | 0.1 | * |
| $\alpha_{10}$ | transition rate from $M_1$ to $M_0$ | $\frac{1}{\text{day}}$ | 0.0001 | * |
| $\alpha_{12}$ | transition rate from $M_1$ to $M_2$ | $\frac{1}{\text{day}}$ | 0 | * |
| $\alpha_{1m}$ | transition rate from $M_1$ to $M_m$ | $\frac{1}{\text{day}}$ | 0.001 | [22] |
| $\alpha_{20}$ | transition rate from $M_2$ to $M_0$ | $\frac{1}{\text{day}}$ | 0.0001 | * |
| $\alpha_{m2}$ | transition rate from $M_m$ to $M_2$ | $\frac{1}{\text{day}}$ | 0.01 | [22] |
| $\beta$ | contribution of tumor to M2-like polarization | — | 2 | * |

In our initial analysis, we explore the influence of changes in the transition rates α_*ij*_ on the steady state values. We start this analysis from a baseline parametrization. For simplicity, we assume for the baseline scenario that transitions from *M*_1_ to *M*_2_ do not happen directly (α_12_ = 0) but go over *M*_0_ or *M*_*m*_, that cells with the mixed phenotype *M*_*m*_ cannot differentiate back to *M*_1_ (*α*_*m*1_ = 0), and that transitions from *M*_2_ to *M*_1_ only happen over *M*_0_ (*α*_21_ = *α*_2*m*_ = 0). Note that increasing these transition rates from 0 to, e.g., 0.001 does not cause any qualitative changes in the results presented in the following (results not shown). In particular, we explore non-zero values for *α*_12_ and its role in bistability in Section 3.1. The values for *α*_1*m*_ and *α*_*m*2_ are taken from [22]. Based on results from a stochastic macrophage polarization model [37], where the transition rate from *M*_0_ to *M*_1_ was larger than that from *M*_0_ to *M*_2_, we fix *α*_01_ = 1 and *α*_02_ = 0.1. We assume the backward differentiation rates to be much smaller and fix *α*_10_ = *α*_20_ = 10^−4^.

To check for multistability, we start simulations with two different initial conditions, one representing high initial tumor volume and one with low initial tumor volume (see Table 2).

**Table 2:** State variables and their initial values

| variable | name | initial value<br>in low tumor | initial value<br>in high tumor |
| --- | --- | --- | --- |
| tumor cells | $T$ | 0.09 | 0.9 |
| naïve macrophages | $M_0$ | 0.1 | 0.1 |
| M1-like macrophages | $M_1$ | 0.1 | 0.1 |
| M2-like macrophages | $M_2$ | 0.1 | 0.1 |
| mixed-phenotype macrophages | $M_m$ | 0.1 | 0.1 |

### 2.2. Analysis on the existence of multistability

Before conducting the numerical analyses provided in later sections below, we first analytically investigate the possibility of multistability by considering the following two theorems.

#### Theorem 1.

*Consider the steady state solution T*^*ss*^ = *of Eq*. (1), *and define* 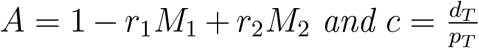. *Then, if 0 < c < A, the non-trivial steady state* 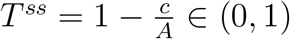.

*Proof*. Setting ^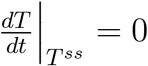^ yields Eq. (6). It implies that a trivial steady state *T*^*ss*^ = 0, and non-trivial solution at 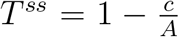 (whose explicit expression is provided in Eq. (7)) also exists. For 0 < c < *A*, it holds 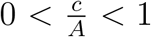, thus the non-trivial *T*^*ss*^ ∈ (0, 1).

Theorem 1 states conditions for which the tumor steady state solution *T*^*ss*^ falls within the meaningful range of (0, 1). We denote the *T*-nullcline of *M*_*i*_(*T*) for i = 0, 1, 2, m, by 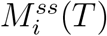, respectively, given that each macrophage population *M*_*i*_(*T*) is a function of the tumor population *T*. Unless necessary, we neglect this fact, however, the dependence of macrophage populations on *T*becomes important in the next Theorem.

#### Theorem 2.

*Let Theorem 1 prevail, and* 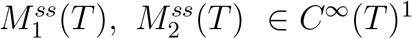 ^1^. *Define H*(*T*) = *p*_*T*_ *T*(1−*T*)*G*(*T*)−d_*T*_ *Twhere* 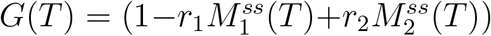. *Then the solution of H*(*T*) *is at least cubic in T*.

*Proof*. Consider the function *H*(*T*) = *p*_*T*_ *T*(1 − *T*)*G*(*T*) − d_*T*_*T* which is (with the term *T*(1 − *T*)) quadratic with respect to *T*. With *G*(*T*) depending on 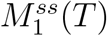 and 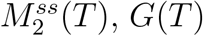 is at least linear in *T*, thus *H*(*T*) is at least cubic in *T*.

Theorem 2 shows that tumor *T*-nullcline is nonlinear (at least cubic) in (0, 1).

#### Corollary 2.1.

*As a consequence of Theorem 1 and Theorem 2, the T-nullcline is S-shaped with up to three roots in the interval* (0, 1).

This implies that the tumor steady state solutions in (0, 1) are either mono-stable or bistable. A tumor steady state is said to be *mono-stable* if there are three intersections 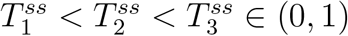 with 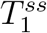 and 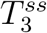 being unstable, and stable states. It is said to be *bi-stable* if there are three intersections 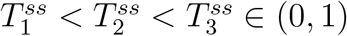 with 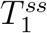 and 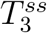 being stable and 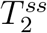 unstable steady states.

To support the analytical results above, a specific formulation of the *T*-nullcline is presented in Appendix B, highlighting key features relevant to bistability.

### 2.3. Methods for model analysis

#### Numerical solution and stability analysis

We use MATLAB’s ode23s solver to solve the ODE system in Eqs. (1)–(5) over 500 time steps. Once the system is solved, we calculate the steady states of the system. To find the steady states as roots of the right hand side of the ODE system, we use the MATLAB function fsolve, which numerically solves a system of nonlinear equations. We then classify the stability of each identified steady state by computing the Jacobian matrix at the steady states and calculating its eigenvalues using the eig function. The Jacobian matrix is calculated analytically (as provided in Appendix C) and its derivation validated using the Symbolic toolbox in MATLAB.

#### Basins of attraction

We calculate the basin of attractions to illustrate how the initial conditions (ICs) evolve towards different steady states (that is, attractors). In the context of our model, it helps visualize macrophage-tumor interactions by plotting the macrophage subpopulations (*M*_0_, *M*_1_, *M*_*m*_, and *M*_2_) against the tumor volume. If an IC cannot be attributed to steady states within the given threshold of 10^−8^, we classify it according to the minimal distance to one of the two tumor volumes (high or low). We calculate of the basin of attraction in the following way:

1. The system of ODEs is solved using the `ode23s` solver for a range of initial conditions. We vary two variables at a time (e.g., tumor vs. *M*_0_, *M*_1_, *M*_2_, or *M*_*m*_) while others are fixed.
2. For each simulation, the trajectory end point (long-term behavior) is compared to known steady states calculated previously.
3. Based on proximity to these steady states, each initial condition is colored, resulting in a 2D plot showing the classification of initial conditions into different basins of attractions.

#### Global Sensitivity Analysis

Global Sensitivity Analysis (GSA) allows the identification of parameters that have the greatest effect on model output. This insight can help identify impactful pathways in the biological system [53]. GSA varies all parameters simultaneously and analyzes the contribution of each (alone, or in groups) to model output. Sobol’s method is one such global method, and is a variance decompostion method, meaning it provides a quantitative measure (termed the sensitivity index) of the contributions of the input to the variance in output. In Sobol’s method, the *first-order sensitivity index* measures the direct contribution of a single input parameter to output variance, while the *total sensitivity index* captures both its direct effect and all interaction effects with other parameters. We compute and present both, and our discussion uses the total sensitivity index to analyze the importance of parameters and pathways for different cases. Sobol’s method is advantageous (for example, over Partial Rank Correlation Coefficient) when there is not a clear monotonic relationship between input and output. We have previously outlined the theory of this method in [27]. Here, we apply GSA to quantify the sensitivity of the tumor volume at steady state with respect to the model parameters. We implemented the GSA using SALib [54], an open-source Python library for sensitivity analysis. We sampled parameter space, varying parameters from baseline values 50% in each direction. The system was solved for each parameter set, steady state values of the tumor volume were recorded, and Sobol’s variance based analysis was conducted using the software.

#### Bifurcation analysis

We generate a bifurcation diagram by plotting steady state end points against a range of chosen bifurcation parameter values (defined through GSA analysis). For the bifurcation plots, we loop through a the range of a specific parameter and solve for each value the ODE system until a steady state is reached. The final values are stored and plotted against the bifurcation parameter values.

All codes used for model analysis are provided to replicate our results presented herein. See Appendix G for more information.

## 3. Simulation Results

### 3.1. Bistable tumor volumes

By varying the values of the transition rates, we observed two qualitatively different steady states: a steady state with *M*_1_ < *M*_2_ and high tumor volume or a steady state with *M*_1_ > *M*_2_ and low tumor volume. A list of the various parameter cases is shown in Table D.5. For example, increasing *α*_1*m*_ or *α*_02_ drives the system from a low-tumor steady state into a high-tumor steady state, thereby passing a region of bistability where two stable steady states co-exist. Figure 2 exemplifies this bifurcation for a change in *α*_1*m*_ from 0.001 (case 1, low tumor volume) over 0.1 (case 2, bistability) to 0.5 (case 3, high tumor volume). A similar transition behaviour can be seen when increasing *α*_02_ from 0.1 (case 1) over 0.5 (case 5, bistability) to 1 (case 6, high tumor volume) or when increasing *α*_12_ from 0 (case 1) over 0.03 (case 8, bistability) to 0.1 (case 9, high tumor volume). In addition, we observed two other parametric configurations that give rise to bistability. Bistability can be reached from case 6 by increasing *α*_20_ from 0.0001 to 0.1 (case 4). Alternatively, bistability can be reached from case 6 by increasing *α*_01_ from 1 to 2 (case 7). A summary over the selected transitions, as described above, can be found in Figure D.6. In what follows in the analysis, we focus on case 2 to discuss the bistability dynamics, unless otherwise indicated.

**Figure 2:**
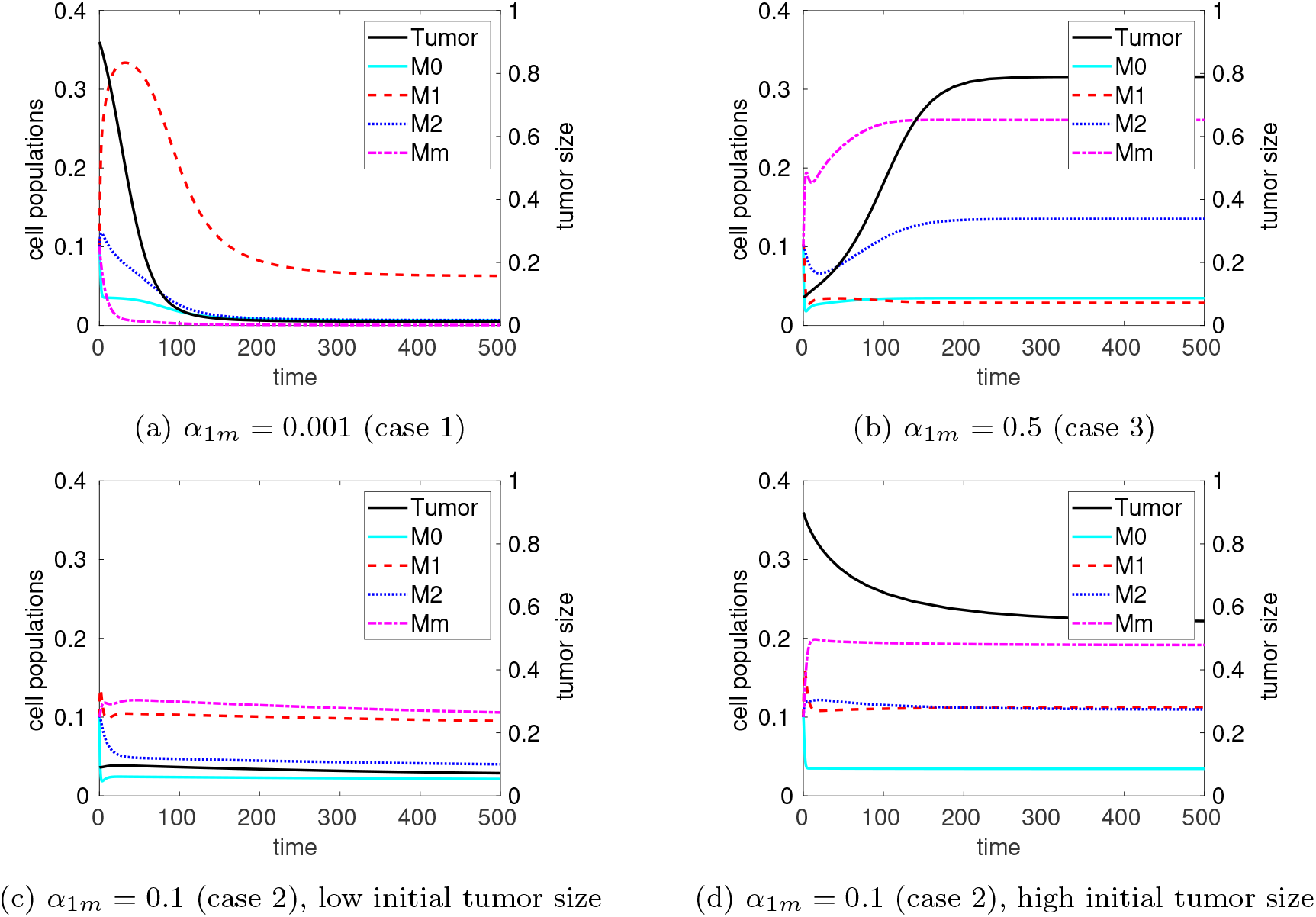
A change in *α*_1*m*_ from 0.001 over 0.1 to 0.5 drives the system from a low-tumor steady state (a) into a high-tumor steady (b) state, thereby passing a region of bistability where two stable steady states co-exist (c,d).

In general, starting from a low-tumor steady state, any increase in transition rates that promote polarization towards *M*_2_ (α_1*m*_, α_*m*2_, α_12_, α_02_) drives the system from a low-tumor steady state over a bistable regimen toward a high-tumor steady state. In contrast, an increase in transition rates that promote *M*_1_ polarization (α_01_ and α_20_) has the opposite effect, i.e., it shifts the system from high tumor steady state over a bistable regimen towards a low-tumor steady state. Note that this type of transition could reflect the treatment with anticancer drugs, which can revert TAM polarization, resulting in increased response to the treatment. Such an example is gemcitabine in pancreatic cancer [55], which induces the polarization of macrophages toward an M1-like phenotype [56]. Similarly, there are also therapies which target Toll-like receptors. For example, Bacillus Calmette-Guerin (a live attenuated strain of bacteria) can reprogram TAMs to become more tumorcidal and increase their inflammatory action. This can again be interpreted as increasing polarization towards M1-like macrophages, which can also be achieved with monoclonal antibodies like anti-PSGL1 [57].

Figure 3 shows the basins of attraction of the two different steady states in the plane spanned by the initial tumor volume and one of the four macrophage sub-populations, *M*_0_, *M*_1_, *M*_2_ and *M*_*m*_, respectively. Depending on the combination between the initial macrophage population and the tumor volume, one of two possible final tumor volumes with either a high or a low volume is reached. It can be seen that the tumor volume in steady state is largely independent from the initial value of *M*_0_, i.e., for higher (or lower) initial tumor volume, the tumor will always end up in the high (or low) steady state. For *M*_1_ and *M*_*m*_, there is a slight dependence of the tumor steady state value on their initial values when the initial tumor volume takes intermediate values around 0.5. In contrast, the tumor volume in steady state highly depends on the initial value of *M*_2_ when the initial tumor volume is below 0.5. In the presence of high initial values for *M*_2_, even a very low initial tumor volume can lead to a high tumor volume in steady state. This illustrates the important influence M2-like macrophages on tumor growth, which has been described in the literature [58].

**Figure 3:**
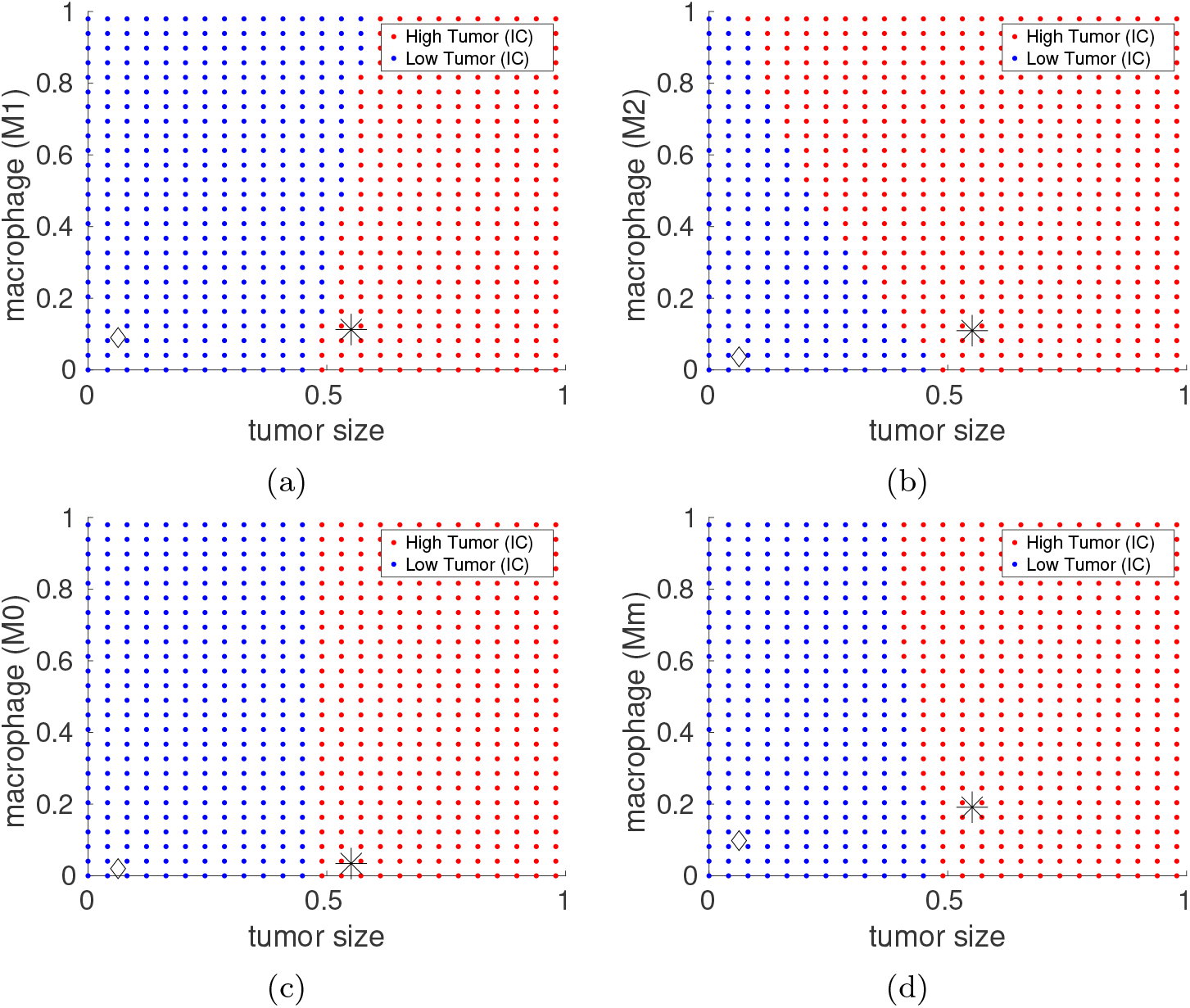
Case 2: Basin of attractions around the following steady states 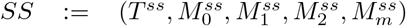 for respectively high (H) and low (L) initial tumor values (*SS*_*H*_ = (0.5511, 0.0343, 0.1126, 0.1095, 0.1914), *SS*_*L*_ = (0.0629, 0.0200, 0.0901, 0.0365, 0.0982)).

### 3.2. Identification of sensitive parameters that drive bistable tumor volumes

We performed GSA for the first three parameter cases 1, 2, and 3 (as shown in Table D.5) to see which parameters are sensitive for low, bistable, and high tumor volume steady state. The GSA results for the three parameter cases are illustrated in Fig. E.7 and Table 3 summarizes the top six most sensitive parameters for each case.

**Table 3:** Top six most sensitive parameters by cases 1, 2, and 3 (in a descending order) with tumor volume as outcome of interest, based on total sensitivity index. Each column refers to low (left), bistable (middle), and high (right) tumor volume cases, respectively.

| case 1 (Low) | case 2 (Bistable) | case 3 (High) |
| --- | --- | --- |
| $r_1$ | $r_1$ | $p_T$ |
| $\alpha_{01}$ | $d_2$ | $d_T$ |
| $d_2$ | $r_2$ | $r_2$ |
| $d_1$ | $\alpha_{1m}$ | $d_2$ |
| $\alpha_{02}$ | $p_T$ | $\beta$ |
| $r_2$ | $\alpha_{01}$ | $K$ |

In case 1, which corresponds to a low tumor volume, the six parameters in decreasing order of total sensitivity indices are *r*_1_, *α*_01_, *d*_2_, *d*_1_, *α*_02_ and *r*_2_. The switching rates from naïve macrophages to either phenotype are identified as having an impact on the variation of the tumor volume in steady state, indicating at low tumor burden, these transitions play a role in the outcome. The apoptosis rates of *M*_1_ and *M*_2_ macrophages also appear on this list (*d*_2_ and *d*_1_). The parameter *r*_1_, which represents the contribution of *M*_1_ cells killing tumor cells, is the parameter with the highest sensitivity index, and it dwarfs the others. The parameter *r*_2_ also appears, but from these results, the impact of *r*_1_ (the contribution of *M*_1_ macrophages killing tumor cells) is more important than the contribution of *M*_2_ macrophages to tumor cell proliferation.

Case 2, which represents a bistable case, the parameters which contribute most to variation in tumor volume are in decreasing order *r*_1_, *d*_2_, *r*_2_, *α*_1*m*_, *p*_*T*_, *α*_01_. Again, *r*_1_ is parameter to which tumor volume is most sensitive. In this case, *α*_1*m*_, appears, indicating in the bistable case, the role of phenotype transition to the mixed phenotype becomes important. Moreover, the parameter *P*_*T*_ appears, which represents the proliferation rate of the tumor. Parameters related to tumor dynamics did not appear in the low tumor volume case, but appear in the bistable case.

When considering case 3 (a high tumor burden case), the six most impactful parameters in decreasing order are *p*_*T*_, *d*_*T*_, *r*_2_, *d*_2_, *β*, and *K*. Here the tumor cell qualities (proliferation rate, death rate) are most important, and phenotype switching rates do not appear. The only parameters relating to macrophages are both related to the *M*_2_ phenotype–its contribution to tumor proliferation and its apoptosis rate. This indicates that the *M*_2_ phenotype is more impactful than the *M*_1_ phenotype in the high tumor volume case. We also see for the first time in the GSA results the parameter K (maximum amount of macrophages relative to tumor volume).

In examining the top six most sensitive parameters across the low, bistable and high tumor volume cases, we find both overlap and divergence in key drivers of system behavior. The parameter *r*_2_ appears in all three cases, indicating that it plays an influential role regardless of the tumor state. Additionally, *d*_2_ and *α*_01_ appear in both the Low and Bistable cases, while d_2_ also appears in the High case, underscoring its relevance in modulating system stability and transitions.

However, the distinct profiles of each case highlight important differences. The low case is characterized by parameters related to the two macrophage phenotypes. The bistable case shares some of this sensitivity but uniquely includes *α*_1*m*_ and *p*_*T*_, indicating that bistability is influenced by additional mechanisms related to transitioning to the mixed phenotype and the tumor’s role in shaping the dynamics. In contrast, the high tumor volume case we see the dominance of tumor-related pathways to the volume at steady state. Together, these patterns reflect a shift in dominant mechanisms as the tumor progresses, with some parameters maintaining relevance across conditions (e.g., *r*_2_), while others exert influence more contextually depending on the tumor state.

### 3.3. Assessing the impact of sensitive parameters on TME through bifurcation analysis

Based on the GSA results in Sec 3.2, we investigate the following bifurcation parameters from the bistable case 2, *r*_1_, *d*_*T*_, *d*_2_, *r*_2_, *p*_*T*_, *β*. However due to the importance of M0 macrophages as unfavourable predictor of overall survival of tumor patients [59], we also investigate the importance of the recruitment of M0 macrophages towards the TME via analysis of *p*_0_ as a bifurcation parameter. We start with demonstrating the impact of parameters whose increase is tumor-suppressing (*r*_1_, *d*_*T*_, *d*_2_, *p*_0_), followed by those that are tumor-promoting (*r*_2_, *p*_*T*_, *β*). Bifurcation diagrams for tumor-suppressive (or -promoting) parameters are shown in Figs. 4 and 5, respectively.

**Figure 4:**
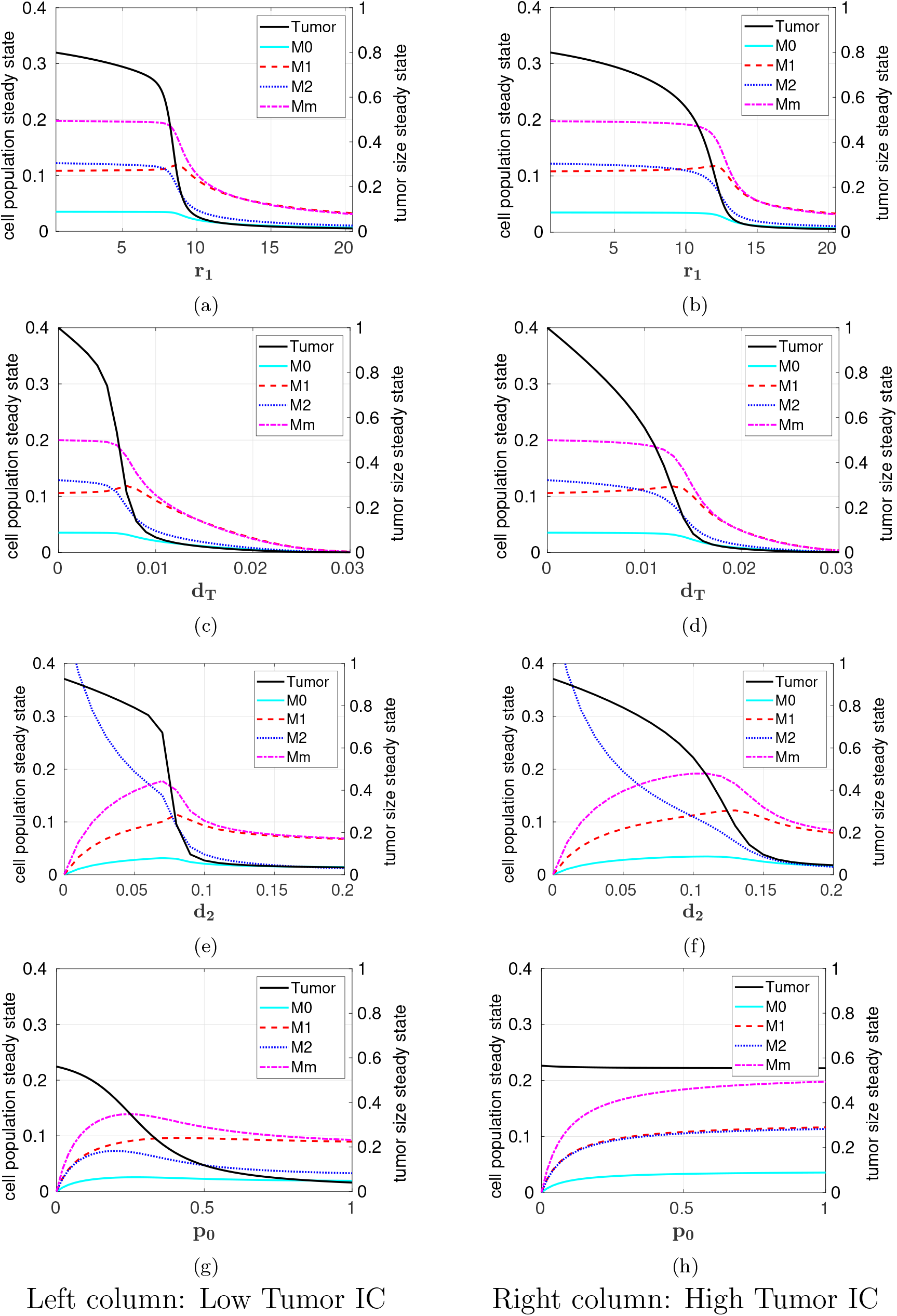
Case 2: Bifurcation analysis for tumor-suppressive parameters, *r*_1_ (top row), *d*_*T*_ (second low), *d*_2_ (third low), and *p*_0_ (bottom row) with low initial tumor volume (left column) and high initial tumor volume (rig1ht7 column).

**Figure 5:**
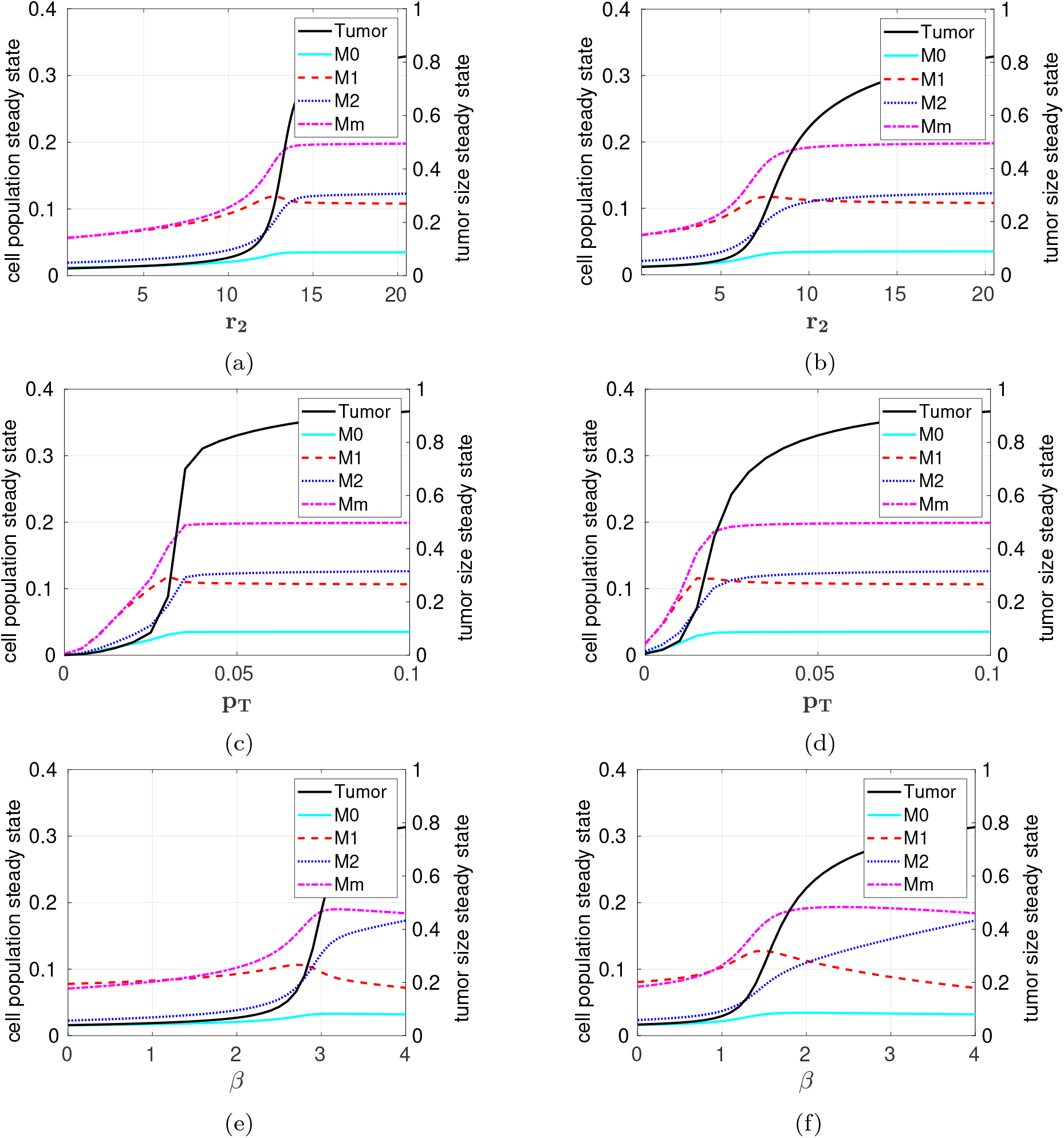
Case 2: Bifurcation analysis for tumor-promoting parameters, *r*_2_ (top row), *p*_*T*_ (middle low), and *β* (bottom row) with low initial tumor volume (left column) and high initial tumor volume (right column).

#### Impact of tumor-suppressive parameters: r_1_, d_T_, d_2_, p_0_

*r*_1_. As shown in Figs. 4a and 4b, increasing *r*_1_ drives the system into a steady state with low tumor burden. The value of *r*_1_ at which this switch happens is larger than the value of *r*_1_ that is needed to keep an initially low tumor volume low. Note that *r*_1_ is the parameter to which the tumor volume is the most sensitive in almost all parameter sets investigated by GSA (except the high tumor burden case). A class of therapeutic medications known as monoclonal antibodies (mAbs)—including rituximab, trastuzumab, cetuximab, and daratumumab, each targeting a specific type of cancer—has been shown to enhance macrophage-mediated phagocytosis and yield positive clinical outcomes [60]. In the context of our model, the parameter *r*_1_ represents the killing rate of tumor cells by M1-like macrophages. Therefore, the therapeutic effects of mAbs can be interpreted as increasing *r*_1_, reflecting enhanced immune-mediated tumor cell clearance. In addition, the molecule CD47, which is normally expressed on healthy cells, acts as a ‘don’t eat me’ signal to macrophages by binding to signal regulatory protein α (SIRPα). Many tumor cells exploit this mechanism by over-expressing CD47, thereby avoiding immune clearance [61]. Immunotherapies that disrupt the SIRPα interaction restore macrophage activity, increasing their phagocytic ability [62]. This therapeutic mechanism can also be captured by an increase of *r*_1_ in our model.

*d*_*T*_. The difference between the low (Fig. 4c) and high initial tumor volume (Fig. 4d) for the parameter d_*T*_ manifests itself in the *d*_*T*_ -threshold value at which the tumor size falls below 0.05. For low initial tumor volume, the threshold is *d*_*T*_ = 0.01 while for high initial tumor volume its threshold is around 0.02. With *d*_*T*_ representing the death rate of tumor cells, a higher apoptosis rate is needed to reduce initially large tumors.

d_2_. Figs. 4e and 4f show a clear correlation between the tumor volume and the size of the *M*_2_ population, which is typically the dominating TAM phenotype in tumors [63, 64]. For low initial tumor size, we observed that an increase in the death rate of M2-like macrophages beyond d_2_ = 0.075 leads to a more abrupt reduction of this population, compared to an initially high tumor volume, where the decay in both populations (M2 and tumor) occurs rather gradually. One can also see that the turning point in both cases is close to the point where the M1/M2 ratio crosses one. This ratio has been identified as a prognostic marker for cancer patients [8].

*p*_0_. The rate *p*_0_ refers to how quickly circulating monocytes are recruited into the TME and differentiated into M0-like macrophages. For high initial tumor volume, a change in *p*_0_ has no impact on the steady state (Fig. 4h). The tumor volume stays high for all positive values of *p*_0_. In contrast, for low initial tumor volume, an increase in *p*_0_ leads to a change from high to low final tumor volume (Fig. 4g). That means, high recruitment of M0-like macrophages (through *p*_0_) helps maintain a low tumor volume through the increase in M1-like polarized TAMs.

This behavior seems to be in contradiction to immunotherapies which target monocyte recruitment. Chemokines and colony stimulating factor 1 (CSF1) are responsible for the recruitment of monocytes to tumors, which then differentiate to become macrophages [65, 66]. Inhibition of the receptor of CSF1 (called CSF1R) has been shown to reduce tumor growth in animal models [67]. Most studies targeting CSF1R are ongoing, so the clinical benefits are not yet reported [68]. Moreover, *p*_0_ is shown to be insensitive in the GSA results and does not greatly contribute to changes in the tumor volume, suggesting that targeting monocyte recruitment might have limited success.

#### Impact of tumor-promoting parameters: r_2_, p_T_, β

*r*_2_. This parameter represents the contribution of M2-like macrophages to the proliferation of tumor cells. Hence, its effect is opposite to this of *r*_1_. Starting from a high initial tumor volume, a lower value of *r*_2_ is needed to change the final tumor volume from high to low (Fig. 5b) compared to the value that is needed to keep an initially small tumor at low volume (Fig. 5a). It is known that M2-like macropahges promote a tumor-supportive environment by inducing tumor cell aggressiveness and proliferation [69]. Cancer cells change their metabolism to be better adapted to the TME [70]. This results in a consecutive metabolic switch of the microenvironment which might be accelerated in larger initial tumor volumes towards a more tumor-supportive environment. An initially low tumor volume might therefore require a larger *r*_2_ to switch the TME than an initially high tumor volume. The same immunotherapies that we discussed in the context of increasing *r*_1_ could be interpreted as decreasing *r*_2_. For example, blocking CD47 signaling can disrupt the tumor-promoting effect of TAMs and potentially enhance the body’s anti-tumor response.

*p*_*T*_. The final tumor volume is particularly sensitive to changes in *p*_*T*_ for values of *p*_*T*_ smaller than 0.04 (Figs. 5c and 5d). Potential therapy options that correspond to a reduction of *p*_*T*_ and thus a reduction of tumor size would include, for example, targeting the glucose uptake in cancer cells [70].

*β*. The parameter *β* describes the strength of influence of the tumor volume on the transition rates α_02_, α_12_, α_1*m*_, α_*m*2_ towards the tumor-supportive macrophage sub-population M2. For a low initial tumor volume, a *β*-value as high as 2.5 is required to switch the M2/M1 sub-population ratio towards an M2 dominating population level. This increase in the M2 population level is also linked to a higher tumor volume in steady state (see Fig. 5e). In contrast, in Fig. 5f it can be observed that for high initial tumor volume and *β* ≤ 1.5, the tumor settles in a low tumor state. However, as soon as *β* increases beyond, so does also the tumor volume and the M2 population size. This indicates that tumor size and M2 population level are directly connected (through *β*). However, the M2/M1 ratio starts to get larger than one at *β* = 2.0, which is earlier than for the low initial tumor volume.

We also analyzed how the region of bistability depends on K, the carrying capacity of the macrophages, and figured out that increasing K shifts the region of bistability towards larger values of *β*. It is meaningful as the impact of the tumor on macrophage polarization needs to be higher if there are more macrophages. Together, the results show that the impact of the tumor towards an M2 dominating environment via *β* needs to be higher for low initial tumor volume than for a high initial tumor volume state. Thus, the variation in *β* has an impact on the M2 population and the M2/M1 ratio.

Therapies that target the macrophage–tumor cell cross-talk include modulation of macrophage polarization, blockade of signaling pathways, and disruption of physical interactions [71].

### 3.4 Impact of sensitive parameters on mono-stable dynamics

For monostable systems, different initial conditions all lead to the same steady state. Still, the steady state values will depend on some of the model parameters, in particular on those that were identified as sensitive by GSA (Table 3). In addition to parameter *β*, we explored variation of the most sensitive parameters *r*_1_ and *p*_*T*_ in mono-stable case 1 and 3 (Fig. F.8).

Interestingly, in case 1, a change in *β* or *p*_*T*_ has almost no influence on the tumor volume in steady state; it remains low despite increasing *β* or *p*_*T*_. In contrast, reducing *r*_1_ leads to an increase in tumor volume at steady state. For case 3, increasing *r*_1_ or decreasing *β* only lead to a very small reduction in tumor volume, whereas decreasing *p*_*T*_ significantly reduces the tumor size.

### 3.5. Biological interpretation of parameters shaping the T-nullcline and driving bistability

We provide a biological interpretation of the model parameters that affect the shape of the *T*-nullcline, thereby governing the system’s potential for bistability. Based on the nullcline shape presented in Appendix B, we observe that the α parameters corresponding to transition rates between different macrophage phenotypes play a critical role in immune system dynamics by regulating the rates at which macrophages switch phenotypes-from unpolarized (M0) to either pro-inflammatory (M1) or anti-inflammatory (M2), and also between M1 and M2.

We also see that the tumor volume modulates macrophage polarization, influencing how quickly M0 becomes M2, M1 becomes mixed or M2 and how M1 and M2 convert into each other through mixed status (see also Eqs. (2)– (5)). It is controlled by the α parameters, which also dictate the polarization rates in response to the tumor volume. The feedback of the tumor population on macrophage landscape is furthermore supported by the parameter *β* ≫ 1, which introduces sigmoidal non-linear relationship.

Both the α values that dictate direct transitions between macrophage phenotypes and those that contribute from the tumor to macrophage feed-back affect how much M1 or M2 is present at steady state. Therewith they determine the strength and direction of the feedback interaction between tumor population and macrophages.

As shown in Eq. (1), macrophages influence tumor volume. So, the states of M1 and M2 directly affect how fast tumor growth is suppressed or promoted. Here we use the parameters *r*_1_, *r*_2_ ≫ 1 to assure that the macrophage influence on tumor growth is strong. This implies that even small changes in α (or *β*) can push the whole system into a different stable state, as we have observed in the above simulations (e.g., Figs. 5e–5f).

## 4. Conclusion

In this study, we present a mathematical model that takes into account how immune cells, specifically different macrophage phenotypes, impact tumor growth and how tumor size impacts the immune system composition. In contrast to previously published models, our model not only includes the classical M1- and M2-like macrophages, but also naïve macrophages and a mixed M1/M2 phenotype as several studies have demonstrated the importance of these subtypes. To our knowledge, there exist only a few previous modeling studies that show bistable dynamics in a cancer-immune context [14, 18, 27, 28]. However, our work develops the first ODE-based population model that explicitly represents three different macrophage phenotypes and a tumor for exhibiting the emergence of bistability via saddle-node bifurcation. Our model results show that bistability makes the system more controllable, responsive to perturbations, and sensitive to immunotherapy.

In fact, we have identified multiple parameters that could act as bifurcation parameters. We explore the effect of altering transition rates (denoted by α_*ij*_) between different macrophage phenotypes on the stability of the system. Though α_*ij*_ are not the most sensitive parameters as shown in GSA results, they appear to be important parameters that enable the system to transition between mono-stable and bistable cases. In a practical setting, it is difficult to alter the transition rates that are promoting *M*_1_ polarization directly by drugs because the underlying molecular mechanisms of these drug actions on the macrophage polarization are complex and involve multiple gene pathways and targets [72]. Drugs that modulate macrophage polarization include the risk of unintended effects on other immune cells, the potential for off-target effects due to broad targeting, and the challenge of achieving precise and localized delivery. In contrast, it seems to be easier to change the composition of the tumor by drugs, e.g. by increasing the death rate of certain cell types, which indirectly affects macrophage polarization. Accordingly, several of the model parameters that were identified as sensitive parameters by GSA are related to such processes (e.g. *d*_*T*_, *d*_2_, *r*_1_, *r*_2_).

The results of our work indicate that when designing an optimized treatment, it depends on where the cancer is in its progression. This is in line with initiatives in precision medicine, and discards the ‘one-size-fits-all’ approach. For low tumor burden, targeting pathways related to macrophages could be impactful. For higher tumor burden, targeting tumor dynamics is preferred. For example, the killing rate of tumor cells by M1-like macrophage (parameter *r*_1_), the apoptosis rate of tumor cells (d_*T*_), and death rate of M2-like macrophages (d_2_) need to be much higher to reduce an initially large tumor compared to the values needed to keep an initially small tumor at a limited size. GSA also revealed that at low tumor volume, it is important to control the differentiation of naïve macrophages towards M1-like or M2-like macrophages (α_01_ or α_02_, respectively) as well as their death rates (d_1_, *d*_2_), whereas at high tumor volume, it is more important to control the dynamics of the tumor volume (*p*_*T*_, *d*_*T*_) and their impact on the macrophages (*β*, K). This reflects that the treatment effect depends on the disease state.

Further analysis and a more detailed model could help to tease out specific thresholds or timing windows for targeted treatment. The bistability observed in our model is a feature of positive feedback loops in its structure, and the multiple intersections of nullclines. Mechanistically, this might be biologically plausible because the immune system fights effectively only until a threshold tumor size is reached [73] at which point the tumor evades the immune system. The tumor-controlled state reflects a balance where the immune system or therapeutic intervention effectively suppresses tumor growth, maintaining the tumor at low or undetectable levels. The tumor-dominated state arises when the tumor escapes immune surveillance or resists treatments. The end game is determined by initial conditions and parameters (which can be altered with therapy), so taking action to drive the system to the tumor-controlled state is the goal.

Despite the richness of our results elucidating the bistable dynamics arising from macrophage-tumor interactions in TME, our work relies on simulation studies only due to the lack of detailed experimental data on bistable tumor behaviors. Experimental setups are often limited in capturing potential alternative outcomes, which hampers reconciling models with data. In fact, though a few modeling studies have attempted to integrate experimental data in investigating the macrophage-tumor microenvironment, the data used is not at the cell-population level [14] or does not show bistability dynamics [22]. To capture biological processes that are difficult to observe experimentally, thus, we apply several computational approaches via numerical simulations, bifurcation and global sensitivity analyses (see Sec. 3).

In addition, macrophages are the only immune cells considered in our model, but there are many others that are known to play an important role (e.g., T cells, B cells, or natural killer cells) in TME [74, 75, 76]. Furthermore, there are additional therapies [68] that we have not discussed in this work, because they would affect parameters that are not sensitive in our model (as informed by GSA results). In our study, the transition rates between different macrophage phenotypes are assumed to be at a constant for analysis simplicity. To alleviate this assumption, a possible direction for future work is to make each transition rate dependent of tumor volume to capture a more biologically realistic process. In addition, many immunotherapy treatments (e.g., mABs [60] and CD47-targeted therapies [61, 62]) associated with increasing phagocytosis affect the same parameter in our model (i.e., *r*_1_ representing contribution of *M*_1_-like macrophages to killing of tumor cells). It is therefore relevant for future work to develop a more refined model to allow different immunotherapies to be investigated at a greater level of details. Moreover, by extending our population-based model for macrophage-tumor interactions, we could further consider combination immunotherapies in TME, known to be an effective strategy from the literature [77, 78].

## CRediT authorship contribution statement

**Hwayeon Ryu**: Conceptualization, Methodology, Conducted numerical simulations, Project administration, Funding acquisition, Writing - original draft, Writing - review & editing. **Susanna Röblitz**: Conceptualization, Formal analysis, Methodology, Conducted numerical simulations, Software, Writing - original draft, Writing - review & editing. **Kamila Larripa**: Conceptualization, Methodology, Conducted the global sensitivity analysis, Software, Writing - original draft, Writing - review & editing. **Anna-Simone Frank**: Conceptualization, Formal analysis, Methodology, Investigation, Conducted the bifurcation analysis, Software, Writing - original draft, Writing - review & editing.

## 6. Declaration of competing interest

The authors declare that they have no known competing financial interests or personal relationships that could have appeared to influence the work reported in this paper.

## 7. Acknowledgment

We thank the American Institute of Mathematics (AIM) for providing a supportive and mathematically rich environment through its Structured Quartet Research Ensembles program, from which this work was first initiated and completed.

## 8. Funding Source

The research of Hwayeon Ryu was funded by National Science Foundation-Association for Women in Mathematic Mentoring Travel Grant through National Science Foundation DMS-#2015440, the Strategic Program for International Research and Education (SPIRE) grant awarded by the University of Bergen, and Elon University Faculty Research & Development Full-Year, Full-Pay Award with Financial Assistance. The research of Susanna Roblitz was funded by Trond Mohn Foundation, Grant No. BFS2017TMT01. Finally, all authors were supported by AIM to travel to the AIM office and meet either in person or virtually each year for 2020–2024.

## Appendix A. Biological pathways notations

Table A.4 that provides a list of notations for biological pathways presented in Fig. 1 and Eqs. (1)–(5).

## Appendix B. Analytical approach to T -nullcline

Setting 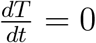 in Eq. (1) leads to

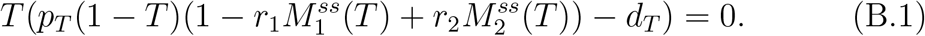

Setting Eqs. (3)–(4) zero individually and solving for 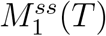 and 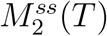 results in

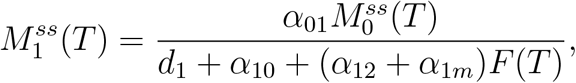

and

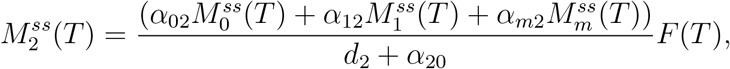

Where 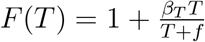. Substituting these expressions into Eq. (B.1) yields

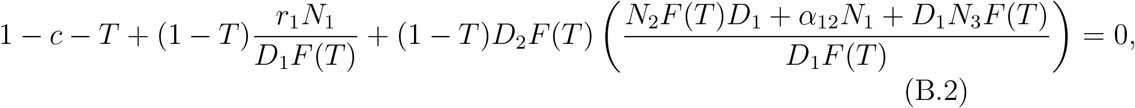

where 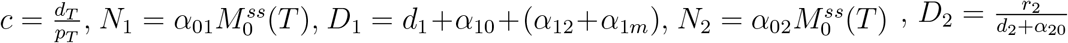 and 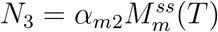.

From last summand in Eq. (B.2), we can see that the *T*-nullcline is a cubic polynomial in *T*. The constants N_1,2,3_ and D_1,2_ depend on the α parameter values, and F(*T*) on *β*_*T*_ parameter value. As can be seen from Eq. (B.2), the tumor-macrophage interactions and feedback, represented by α and *β*, influence the shape of *T*-nullcline and thus the multistable state of the system.

## Appendix C. Derivation of Jacobian equations

Jacobian equations for our model equations, (1)–(5), are derived under the following notations: 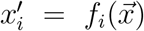 for i = 1, 2, 3, 4, 5, where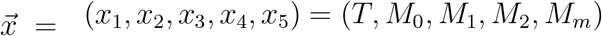 and 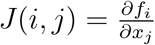.

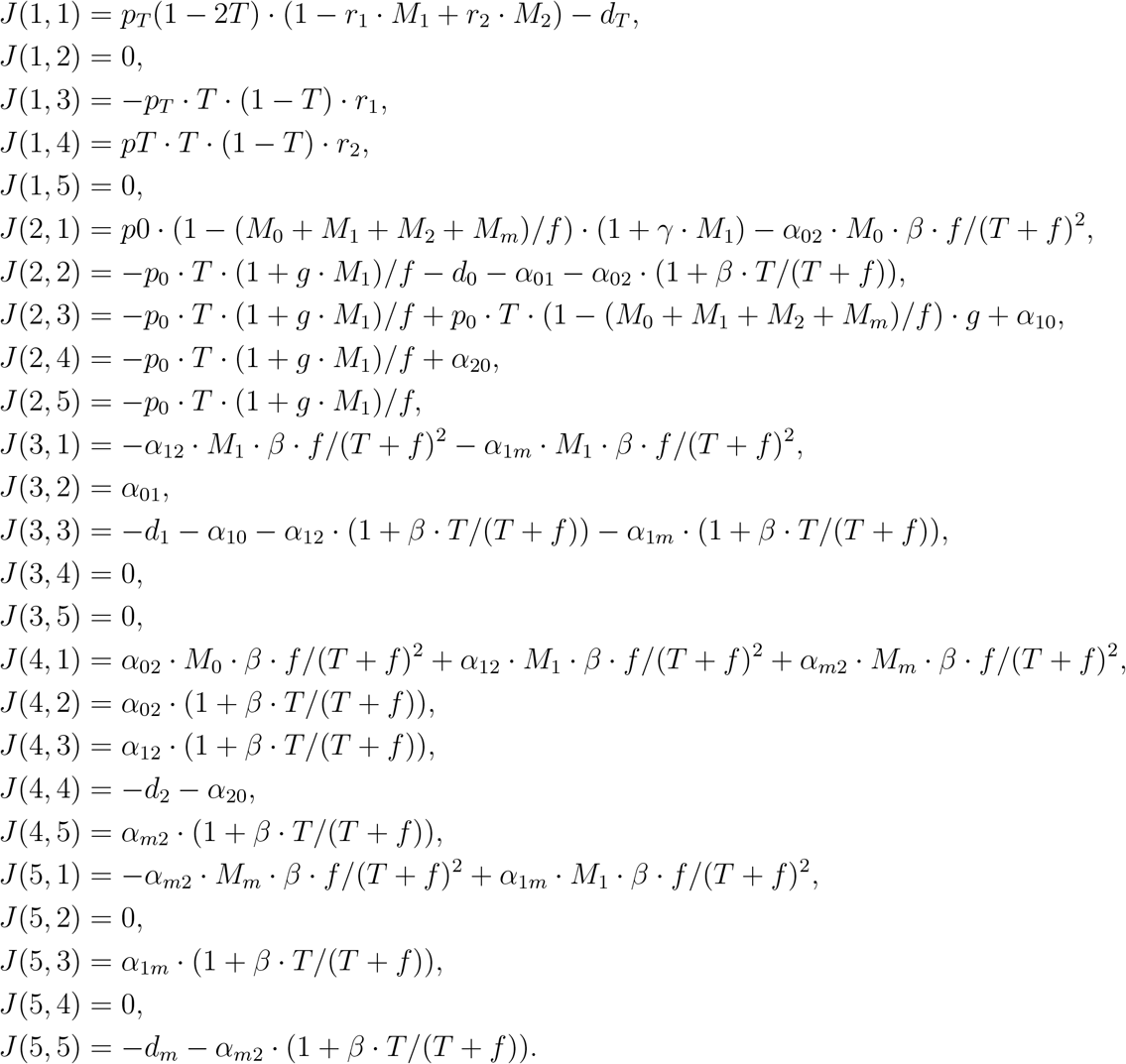

## Appendix D. Parameter cases

Table D.5 summarizes the list of all cases considered to demonstrate parameter variations leading to the different system’s steady states (i.e., with different tumor volumes). The corresponding flow chart of selected case transitions is shown in Figure D.6 from low tumor volume (red; left column), over bistability (green; middle column) to high tumor volume (blue; right column) by altering selected transition rates α_*ij*_.

**Figure D.6:**
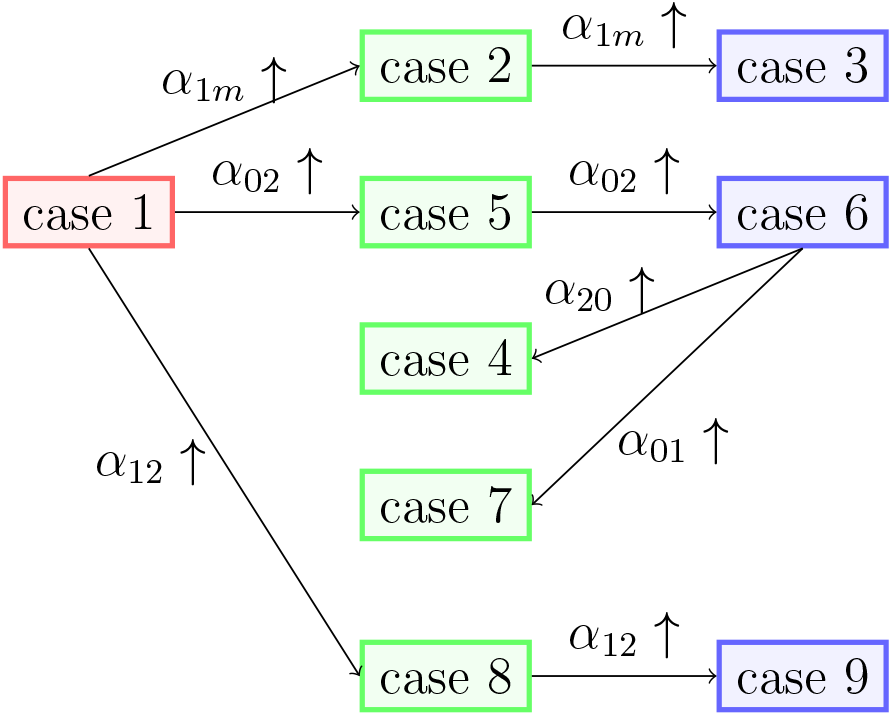
Flow chart of case transitions from low tumor volume (red; left column), over bistability (green; middle column) to high tumor volume (blue; right column), by increasing selected parameters *α*_1*m*_ (top transition), *α*_02_ (second transition), and *α*_12_ (bottom transition). Note that all parameter values used for each case are provided in Table D.5. Two additional bistability cases, case 4 and case 7, can be obtained from case 6 by increasing parameters that promote tumor suppression, *α*_20_ and *α*_01_, respectively.

## Appendix E. GSA plots

Figure E.7 shows the results from a Global Sensitivity Analysis (in Sec. 3.2) for parameter cases 1, 2, and 3 (as shown in Table D.5).

## Appendix F. Figures for mono-stable bifurcation analyses of most sensitive parameter and β

Figure F.8 shows the impact of *β* and other sensitive parameters (*r*_1_ and *p*_*T*_) on the steady-state tumor volume in mono-stable cases 1 and 3.

## Appendix G. Code

Codes for bifurcation diagrams and graphs of population dynamics (shown in Section 3) will be published in GitHub upon the acceptance of the article. Also, Python code used to create the GSA figures (Fig. E.7) is available on the Github repository: https://github.com/Larripa/Macrophage-Tumor-Population-Model-GSA.

**Figure E.7:**
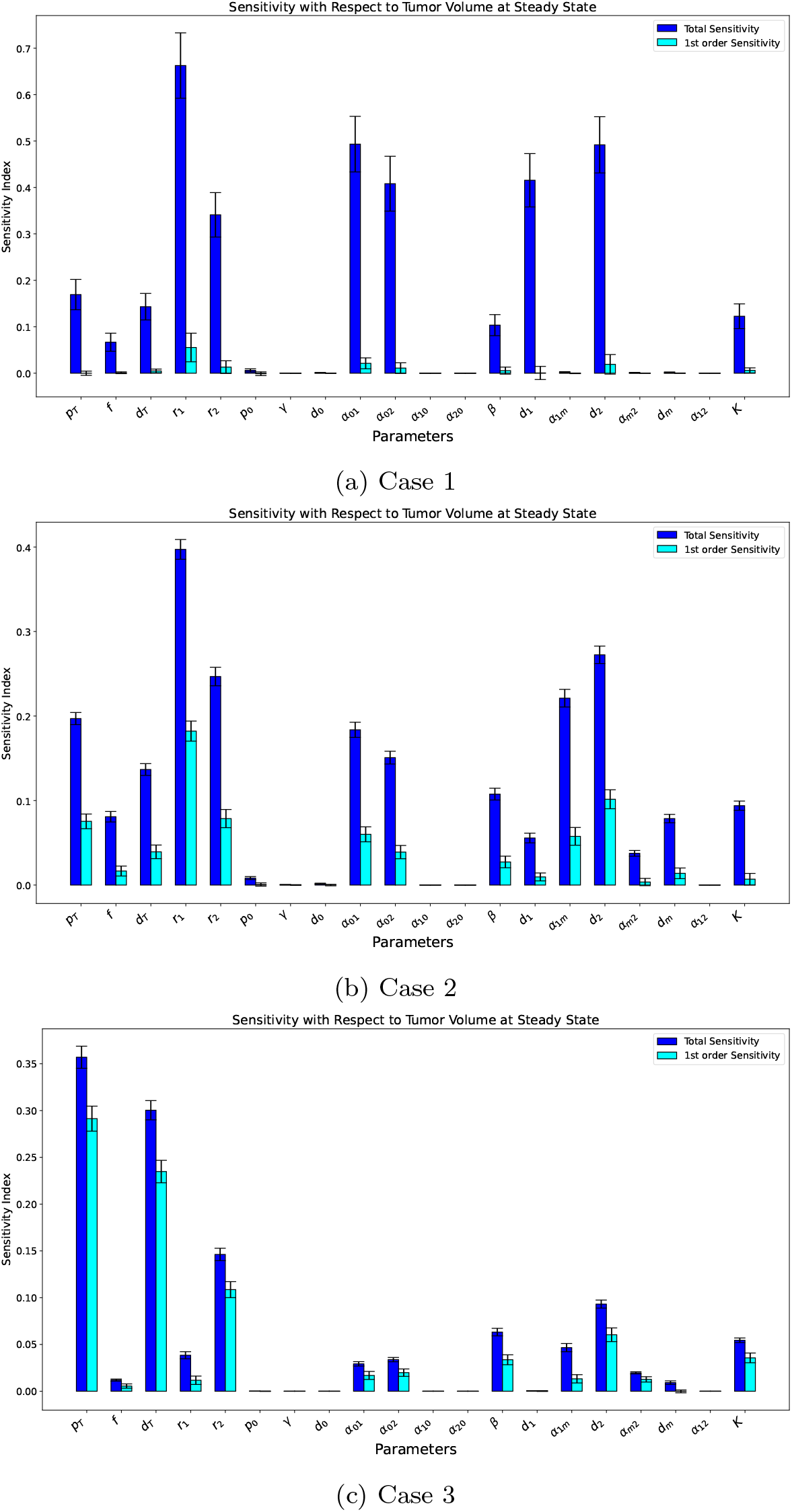
GSA results for three parameter sets 1 (top), 2 (middle) and 3 (bottom), respectively. The figures show the Sobol Sensitivity Indices (first order and total sensitivity) where the outcome of interest is tumor volume at steady state. The GSA used baseline parameter values presented in Tables 1 and D.5 and varied them 50% in each direction. Parameters listed with a value of zero were simulated as 10^−8^.

**Figure F.8:**
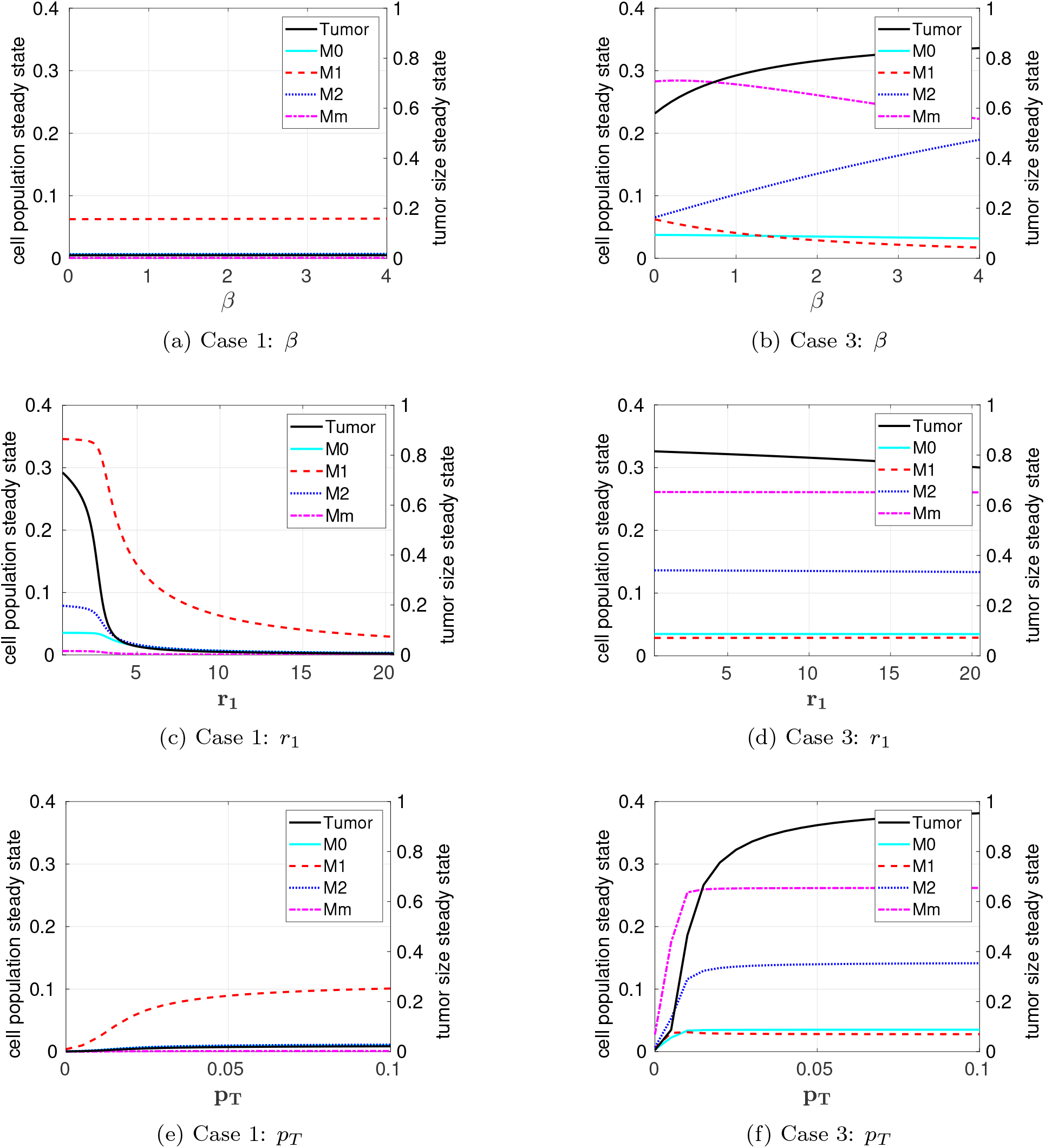
Impact of *β* and other selected sensitive parameters (*r*_1_ and *p*_*T*_) on the steady-state tumor volume in mono-stable cases 1 and 3.

**Table A.4:** Notation of biological pathways presented in Fig. 1

| Pathway | Description |
| --- | --- |
| $a$ | Logistic growth of tumor cell number, which is inhibited by the presence of M1-like macrophages but stimulated by the presence of M2-like macrophages |
| $b$ | Natural death rate of tumor cells, including both necrosis and apoptosis |
| $c$ | Monocyte recruitment |
| $d$ | Natural death rate of naïve macrophages |
| $e$ | Transition of Naïve macrophages to M1-like phenotype |
| $f$ | Transition of naïve macrophages to an M2-like phenotype |
| $g$ | Transition of M1-like macrophages to a naïve phenotype |
| $h$ | Transition of M2-like macrophages to a naïve phenotype |
| $i$ | Natural death rate of M1-like macrophages |
| $j$ | Transition of M1-like macrophages to M2-like macrophages |
| $k$ | Transition of M1-like macrophages to mixed phenotype macrophages |
| $l$ | Natural death rate of M2-like macrophages |
| $m$ | Transition of Mixed-phenotype macrophages to M2-like macrophages |
| $n$ | Natural death rate of mixed phenotype ($M_m$) macrophages |

**Table D.5:** Parameter variations that change the system’s steady states between low tumor volume, high tumor volume, and bistable regimens.

| case | steady state | $\alpha_{02}$ | $\alpha_{1m}$ | $\alpha_{01}$ | $\alpha_{20}$ | $\alpha_{12}$ |
| --- | --- | --- | --- | --- | --- | --- |
| 1 | low | 0.1 | 0.001 | 1 | 0.0001 | 0 |
| 2 | bistable | 0.1 | 0.1 | 1 | 0.0001 | 0 |
| 3 | high | 0.1 | 0.5 | 1 | 0.0001 | 0 |
| 4 | bistable | 1 | 0.001 | 1 | 0.1 | 0 |
| 5 | bistable | 0.5 | 0.001 | 1 | 0.0001 | 0 |
| 6 | high | 1 | 0.001 | 1 | 0.0001 | 0 |
| 7 | bistable | 1 | 0.001 | 2 | 0.0001 | 0 |
| 8 | bistable | 0.1 | 0.001 | 1 | 0.0001 | 0.03 |
| 9 | high | 0.1 | 0.001 | 1 | 0.0001 | 0.1 |

## Footnotes

1 The class of functions that have infinitely many continuous derivatives with respect to *T*.

## Notes

### Competing Interest Statement

The authors have declared no competing interest.

